# Extended DNA damage-induced G2/M arrests lead to aberrant mitoses and cell death due to excessive accumulation of securin

**DOI:** 10.1101/2025.07.21.665839

**Authors:** Anton Dubenko, Marah Jnied, Oleksii Kotenko, Svetlana Makovets

## Abstract

The stochastic nature of DNA damage dictates various scenarios of stress survival among different cells. We used time-lapse microscopy to perform single-cell level analyses of yeast populations with double-stranded DNA breaks and nonfunctional telomeres. Activation of the DNA damage signalling resulted in a broad distribution in the duration of G2/M arrests. Strikingly, the longer arrests correlated with aberrant mitoses caused by mis-coordination of nuclear division and cytokinesis, leading to cell death. Chk1-dependent phosphorylation of securin, an inhibitor of sister chromatid separation in mitosis, was responsible for this phenomenon. Securin progressively accumulated during G2/M arrests, leading to grossly delayed or missed nuclear divisions. This phenotype could be suppressed by slowing down the progression of mitotic exit via *LTE1* deletion. Lowering securin levels also partially supressed the DNA damage-induced aberrant mitoses but resulted in an increase in aneuploidy during normal growth. Our results demonstrate that in cells taking too long to complete DNA repair, the DNA damage checkpoint promotes aberrant mitoses and cell death, thereby eliminating cells with a higher chance of genomic instability. This mechanism in microbial populations might parallel the senescence program in mammals where long cell cycle arrests become irreversible and also lead to cell death.

**IMPORTANT:**

- Manuscripts submitted to Review Commons are peer reviewed in a journal-agnostic way.
- Upon transfer of the peer reviewed preprint to a journal, the referee reports will be available in full to the handling editor.
- The identity of the referees will NOT be communicated to the authors unless the reviewers choose to sign their report.
- The identity of the referee will be confidentially disclosed to any a filiate journals to which the manuscript is transferred.

**GUIDELINES:**

- For reviewers: https://www.reviewcommons.org/reviewers
- For authors: https://www.reviewcommons.org/authors

**CONTACT:** The Review Commons office can be contacted directly at

## INTRODUCTION

DNA damage signalling is highly conserved in eukaryotes (Melo and Toczyski 2002). It is triggered in response to DNA breaks and stalled replication forks. The DNA damage sensor kinases Mec1 and Tel1 in yeast and their homologues ATM and ATR in mammals, are recruited to such structures, triggering a protein phosphorylation cascade, where Mrc1 and Rad9 mediators facilitate activation of the downstream effector kinases Chk1 and Rad53 (CHK1 and CHK2 in mammals). The effector kinases phosphorylate multiple proteins required to stop cell cycle progression and activate DNA repair.

In yeast, the Pds1 protein called securin is one such target for Chk1 kinase (Sanchez et al. 1999; Wang et al. 2001). Securin is an inhibitor of separase Esp1 (Ciosk et al. 1998), which catalyses proteolytic cleavage of cohesin to allow sister chromatid separation in mitosis (Uhlmann et al. 1999). The Mec1-Rad9-Chk1-dependent phosphorylation of Pds1 stabilises securin and delays sister chromatid separation (Cohen-Fix and Koshland 1997; Sanchez et al. 1999; Wang et al. 2001). This regulatory pathway plays an important role in holding cells with DNA damage in G2/M while DNA repair takes place. At the metaphase-to-anaphase transition, Pds1 is dephosphorylated and degraded by the proteasome (Cohen-Fix and Koshland 1997; Agarwal et al. 2003). As a result, separase is no longer inhibited and the sister chromatids can be segregated to daughter cells.

Nonfunctional telomeres resemble DNA breaks (Sandell and Zakian 1993). If they are too short to recruit enough telomere-binding proteins, which shield the chromosomal DNA ends, the DNA damage sensor kinases recognise them as broken DNA and activate the damage signalling pathway (Enomoto et al. 2002; d’Adda di Fagagna et al. 2003). Telomeres are extended by the enzyme telomerase, which compensates for the telomere shortening during replication (Greider and Blackburn 1985). Either an insufficiency or a complete loss of telomerase function results in gradual telomere shortening after every replication cycle (Lundblad and Szostak 1989; Harley et al. 1990; Lundblad and Blackburn 1990; Yu et al. 1990). This process is called replicative senescence and eventually leads to critically short nonfunctional telomeres, which activate DNA damage signalling and cell cycle arrest.

In humans, the expression of the catalytic subunit of telomerase hTERT is transcriptionally downregulated during embryonic development (Sharma et al. 1995; Holt et al. 1996; Roake and Artandi 2020). The resultant telomerase insufficiency (TI) is important for inhibiting carcinogenesis by limiting cell proliferation through replicative senescence. However, telomerase becomes re-activated in 85-90% of cancers and the remaining 10-15% switch to ALT (Alternative Lengthening of Telomeres), a recombination-dependent mechanism for telomere maintenance (Artandi and DePinho 2010; Cesare and Reddel 2010). Similarly, a loss of the telomerase catalytic subunit Est2 in yeast leads to replicative senescence, activation of the DNA damage signalling and G2/M arrest, followed by cell death of the majority of cell population (Lendvay et al. 1996). However, rare survivors emerge. They are similar to the ALT cancer cells in using recombination for telomere maintenance (Lundblad and Blackburn 1993; Lendvay et al. 1996). Therefore, the telomere dynamics and its interplay with the DNA damage signalling pathway share a lot of similarities in yeast and human cells.

Cell populations experiencing either DNA breaks or telomere maintenance problems are heterogeneous because of the probabilistic nature of DNA damage and telomere dynamics: not every cell in the population experiences the same number of breaks or nonfunctional telomeres in every cell cycle. In addition, the progress of the repair could be different too. The combination of both variabilities would then result in variable, highly individualised cellular response and consequent fate. Often, the dynamic of the DNA damage response is analysed at a population level, thus producing results that average potentially an extremely wide variety of responses and outcomes at the level of individual cells. This blunts the sensitivity of our measurements: do all cells in a population experience the same delay in response to damage? Do they have the same outcome of surviving the DNA damage stress? To answer these questions, we used time-lapse microscopy in combination with fluorescently-tagged proteins to investigate budding yeast with telomerase insufficiency, telomerase deficiency and phleomycin-induced DSBs at the single-cell level. In all these settings, we observed a variety of different cell behaviours, from no cell cycle arrest in some cells to a very long time spent in G2 by others. Importantly, the longer cell cycle arrests often resulted in aberrant mitoses leading to cell death, thereby eliminating the cells with hard-to-repair or extensive DNA damage from the populations. Co-visualisation of several cellular components combined with genetic analyses allowed us to gain molecular insights into the mechanisms of this pathway, where accumulation of Pds1 during G2/M arrests operates as a stopwatch differentiating the cells with shorter and longer arrests.

## RESULTS

### Yeast telomerase insufficiency (TI) model system

In a haploid state, the budding yeast *Saccharomyces cerevisiae* carry 16 chromosomes and 32 telomeres. All the telomeres contain ∼350 bps of (TG_1-3_)_n_ telomeric repeats, as well as repetitive X-elements in the sub-telomeric regions (Figure 1A). About two thirds of all the telomeres also contain one or more Y’ repeats inserted between (TG_1-3_)_n_ and X-elements (Wellinger and Zakian 2012). The telomeres with Y’ repeats are called Y’-telomeres and the rest are X-telomeres (Figure 1A). Upon telomerase loss, the (TG_1-3_)_n_ telomeric sequences shorten with every replication cycle and eventually most cells in the population die due to nonfunctional telomeres. However, a small fraction of cells survives by switching to one of the two recombination-dependent mechanisms of telomere maintenance. In Type I survivors, all the X-telomeres acquire Y’ repeats and homologous recombination involving Y’s maintain short but stable (TG_1-3_)_n_ telomeric tracts, while amplifying Y’ repeats. In Type II survivors, recombination operates on the (TG_1-3_)_n_ telomeric tracts which become very long as recombination amplifies primarily these repeats.

**Figure 1.**
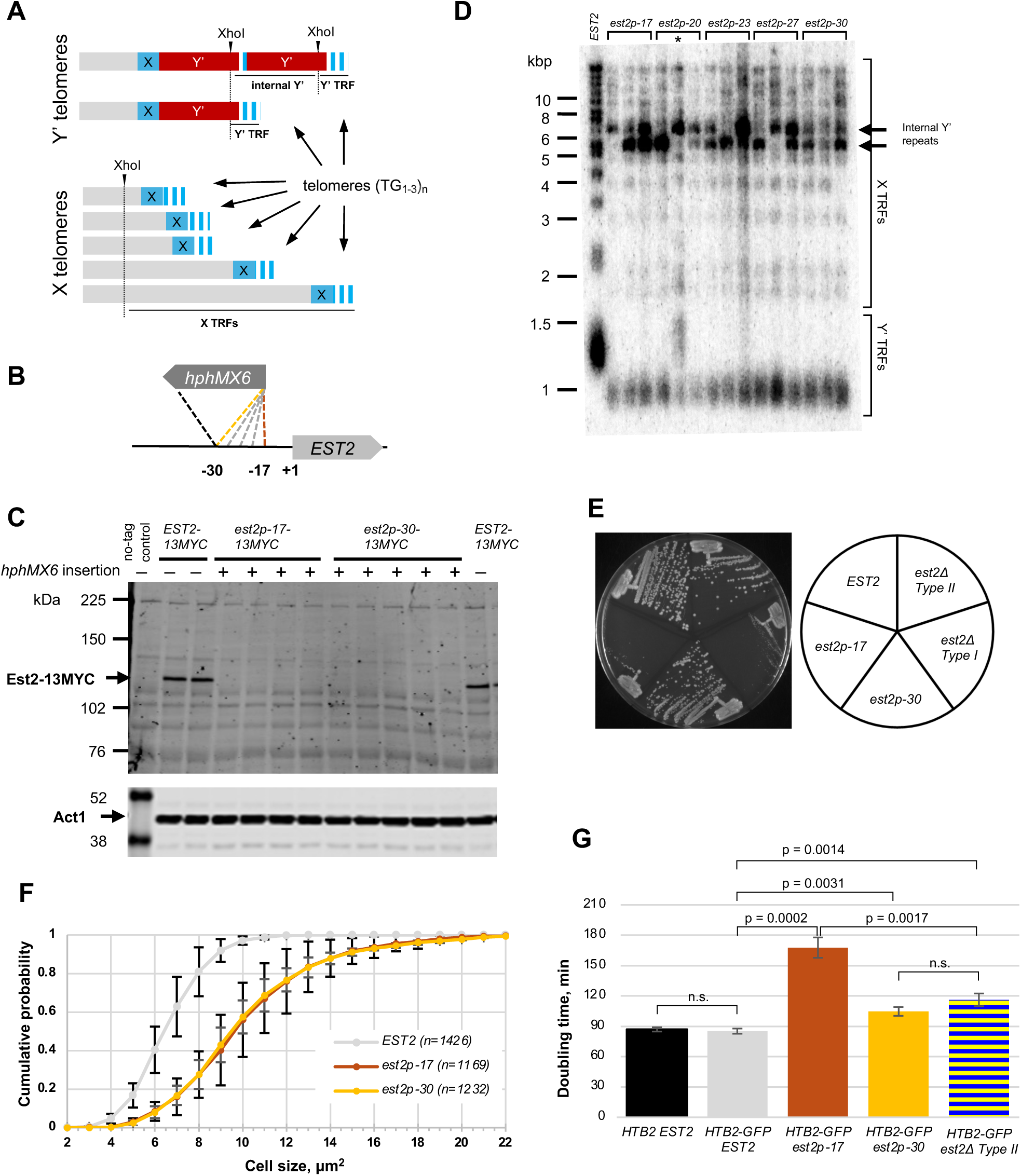
Construction of yeast strains with TI. (A) Schematic of yeast telomeres with the subtelomeric repetitive elements shown. All telomeres contain subtelomeric X-elements (in blue) and the actual telomeric (TG_1-3_)_n_ repeats (blue-white stripes). Y’ telomeres also include one or more Y’ repeats (in dark red) separated by short stretches of (TG_1-3_)_n_, while the X-telomeres lack Y’s. A restriction digest of genomic DNA with XhoI (dotted lines, cut sites shown by arrows) generates terminal restriction fragments (TRFs), which run on a gel as smears due to the variable length of (TG_1-3_)_n_. The XhoI site in Y’ generates similar TRFs in all the Y’ telomeres. Because XhoI sites are outside of the repetitive elements in X-telomeres, each X telomere generates a unique TRF. TRFs from X-telomeres run as multiple smeary bands, all above the Y’ TRFs as shown in panel D, where TRFs are visualized on a Southern blot using a probe hybridising to (TG_1-3_)_n_. (B) Schematic for mutating *EST2* promoter via integration of *hphMX6* upstream of the coding region. The position of the first coding nucleotide (A in the ATG) is marked as +1. The positions of the *hphMX6* insertions in the promoter region are indicated relative to the *EST2* coding region. (C) Western blot analysis of Est2-13MYC levels in cells with different *hphMX6* promoter insertions. The bottom panel shows Act1 blot as a control for sample loading. The same protein samples were loading in in corresponding lanes for both blots, except in the left most lane, where the no myc-tag control was omitted and the marker lane is shown instead on the Act1 blot image. (D) Telomere length and structure analysis of *est2p* mutants by Southern blotting. 3 independent clones for each genotype were analysed after 10 passages (∼200 generations). Parental strain with the native *EST2* promoter is shown in the first lane. Oligonucleotide hybridising to the telomeric (TG_1-3_)_n_ sequence was used as a probe to visualise telomeres. One of the clones (marked with *) generated Type II survivors. (E) Comparative analysis of cell growth on YPD plates. The set of isogenic strains includes the parental *EST2* yeast, the TI mutants *est2p-17* and *est2p-30* with equilibrated telomeres, as well as telomerase-deficient Type I and Type II survivors. Cells were streaked from fresh patches onto YPD plate and incubated for 2 days at 30°C before the image was taken. (F) Cumulative distribution of the average cell size among cell populations with the normal amount of telomerase and with TI. (G) Population doubling time of TI and telomerase-deficient Type II survivors in the *HTB2-GFP* strain background. Averages (shown below each bar) of at least 3 biological repeats are plotted. The error bars represent st.dev. Here and in other plots below, P-values for different pairwise comparisons are shown above the corresponding square brackets indicating the pairs. n.s. stands for non-significant, i.e. p > 0.05 by Student’s two-sample two-tailed t-test.

Most work on how yeast deal with nonfunctional telomeres has been done using cells with a complete telomerase loss due to a deletion of a telomerase gene. To engineer a model system to study TI in yeast, we mimicked the effects of the developmentally programmed downregulation of hTERT in humans by weakening the *EST2* promoter. A gene cassette encoding a Hygromycin resistance gene (*hphMX6*) was inserted into the promoter in the orientation opposite to that of *EST2* (Figure 1B). The terminator end of the cassette was integrated 30 bp away from the *EST2* translation start site (the −30 locus), while the position of the *hphMX6* promoter end varied in different mutants, from −30 to −17, to create a set of *est2p*-alleles (for *<u>p</u>*romoter). Analysis of *est2p-17-13MYC* and *est2p-30-13MYC* cells by western blotting showed a decrease in Est2-13MYC protein to undetectable levels (Figure 1C). Multiple isolates from independent integration events were passaged 10 times on YPD plates and their telomeres were analysed by Southern blotting (Figure 1D). Although the *est2p* mutants showed a noticeable amplification of Y’ repeats characteristic of telomerase-deficient Type I survivors, in most isolates the X-telomeres resembled those of the telomerase-positive parental strain, but equilibrated at a much shorter length, indicating telomerase-dependent telomere maintenance. This was consistent with the lack of a viability crisis normally present during *est2Δ* strain passaging. Although the *est2p-30* cells grew significantly better on agar plates than the *est2p-17* mutants (Figure 1E), both promoter mutations led to an increase in the fraction of large cells in the populations (Figure 1F). This is consistent with a frequent activation of the DNA damage checkpoint when the arrested cells experience excessive growth during prolonged G2.

### TI leads to a significant increase in aberrant mitoses, accompanied by loss of nucleolus and cell death

To study TI at the single-cell level using time-lapse microscopy, we introduced the *est2p-17* and *est2p-30* alleles into the strain background with the GFP-tagged histone H2B (Huh et al. 2003; Crane et al. 2019), passaged cells on YPD plates 10-15 times to establish cultures with equilibrated short telomeres and confirmed them retaining their original telomere structure by Southern blotting (short telomeres, but X-telomeres retained), as described above. As expected, both *est2p-17* and *est2p-30* mutations increased the *HTB2-GFP* population doubling time in YPD broth (Figure 1G), with the former causing a more pronounced effect, consistent with the growth defect observed on agar plates (Figure 1E). The *HTB2-GFP* allele did not cause any growth defect in the *EST2* background (Figure 1G). The *HTB2-GFP EST2* strain is referred to as wild-type hereafter.

We followed cell divisions over an extended period of time (4-8 h) using time-lapse microscopy, with an image taken every 5 min. Most of the *EST2* cells completed mitoses normally. As expected, they first moved their nuclei to the bud neck and underwent nuclear divisions 5-10 minutes later (1 or 2 frames). We called these mitotic events Normal Divisions (see ND in Figure 2A, top left and Figure S1A, top panel). In contrast, a large fraction of cells with TI experienced cell cycle arrests characterised by extended time spent with the nuclei held at the bud neck. In addition, they showed a distinctly abnormal nuclear morphology, where the majority of the genome was transferred into the bud while a thread-shaped Htb2-GFP signal corresponding to the nucleolus (see below) remained in the mother cell (Figure 2A, top right.) There were 3 distinct outcomes following the arrests. Each was categorised to a specific class, 1 through 3. Class 1 included arrested cells which eventually managed to undergo seemingly symmetrical nuclear divisions and both mother and daughter cells were able to proceed through the subsequent cell cycles (Figure 2A and S1A, Class 1). Class 2 cells remained arrested at the time of imaging termination (Figure 2A, Class 2). Thus, these were the G2-arrested cells with unclear outcomes of their mitoses, either normal (Class 1) or aberrant (Class 3), at the end of imaging. Keeping track of these events was important for the overall analysis of the populations, as the Class 2 events reflected the overall fraction of cells undergoing cell cycle arrest, as well as the duration of the arrests. Class 3 included cells that went through aberrant mitoses with clearly observed nuclear missegregations (Figure 2A, Class 3). These aberrant mitoses were the main focus of this study.

**Figure 2.**
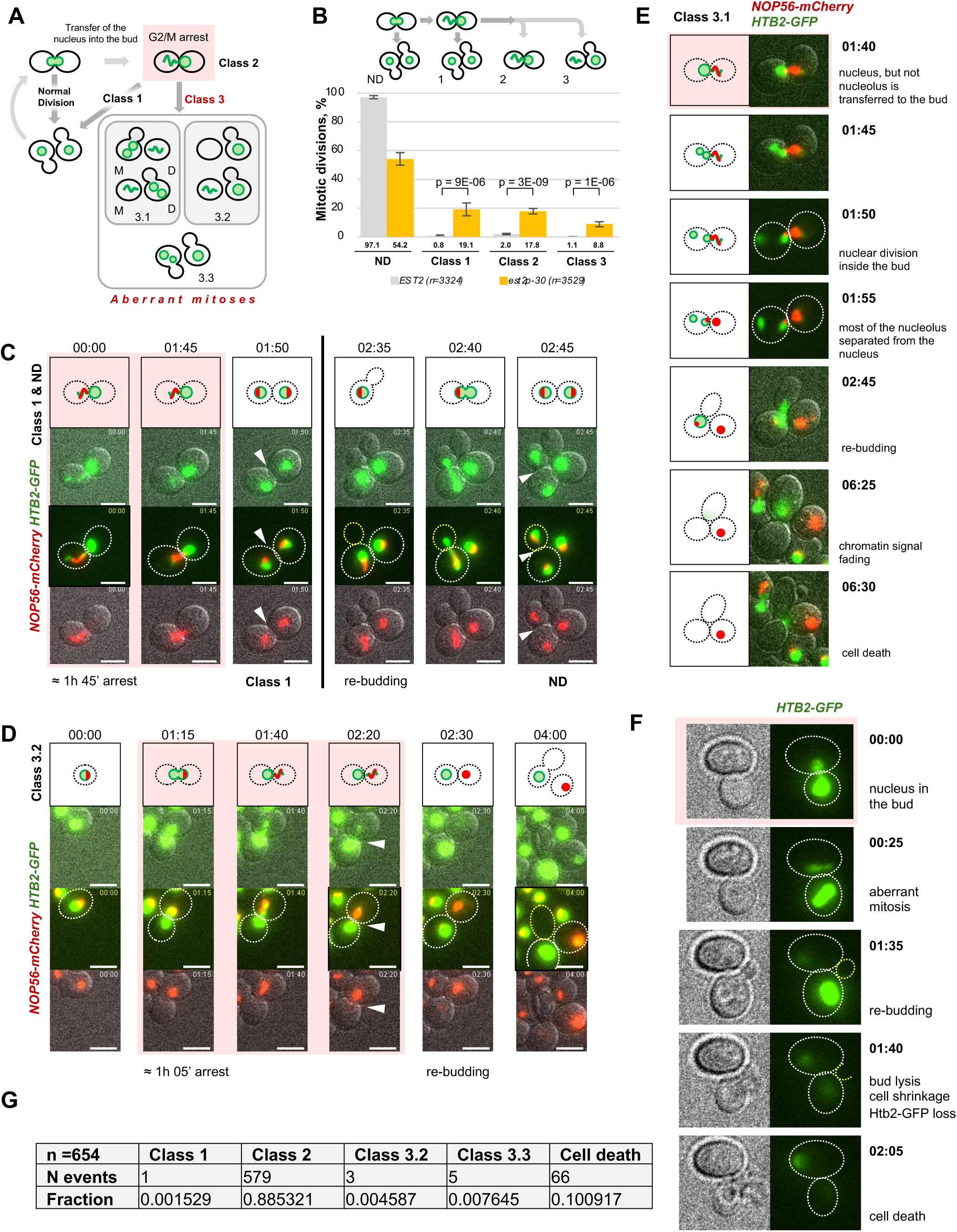
TI leads to a significant increase in aberrant mitoses, accompanied by loss of nucleolus and resulting in cell death. (A) Schematic describing the classification of mitotic divisions in time-lapse microscopy experiments. See text for explanations. (B) TI in *est2p-30* leads to an increase in aberrant mitoses. Averages (shown below each bar) of at least 3 biological repeats are plotted for each genotype. The error bars represent st.dev. A total number of mitotic events (n) scored for each genotype in all the repeats is stated in the legend. (C-E) Representative examples of mitotic events in cells with TI: Class 1 and ND in C; Class 3.2 in D and Class 3.1 in E, with co-visualized nucleus (Htb2-GFP) and nucleolus (Nop56-mCherry). Consecutive images of the same field during time-lapse microscopy are shown in each set. For each image, a schematic depicting nuclear (in green) and nucleolar (in red) dynamics are shown above (C-D) or next to the image (E). The time (hours: minutes) shown above each schematic (C-D) or on the left of each image (E) is relative to the start of the imaging. This time was used to estimate the length of cell cycle arrests (shown by pink background) and other events. Notice that in panel C, Class 1 and ND events come from imaging two consecutive divisions of the same cell. Arrows point at nuclear divisions. Here and in the figures below, white cell outlines are used for the initial cell divisions and yellow outlines for the later emerging buds (re-budding) and the cells derived from them. (F) A representative example of cell death with clearly observed bud lysis following an aberrant mitosis. (G) Analysis of cell fate following aberrant mitoses. 654 *est2p-30* cells, which underwent aberrant mitoses, were analysed in the following cell cycle and the observed results were summarised in the table.

Depending on the observed nuclear dynamics during aberrant mitoses, we defined 3 subclasses in Class 3, from 3.1 to 3.3 (Figure 2A and S1A). Class 3.1 included mitotic events with either a completed or an attempted and reversed nuclear division within a single cell compartment (normally inside the bud) resulting in one cell retaining the majority of the DNA in the form of a single nucleus or two nuclei, while the other cell inherited very little or no visible DNA. Class 3.2 divisions were characterised by a completely skipped nuclear division resulting in a cell with the majority of the duplicated genome, predominantly ending up in daughter cells, and an anuclear cell. In both Class 3.1 and 3.2, the genome-containing daughters re-budded, arrested in the next cell cycle again and in a vast majority of cases died, whereas the mothers remained cadavers with little DNA content. Class 3.3 included any cell divisions resulting in cells with more than two nuclei generated during a single mitosis.

Analysis of the frequencies of different mitotic events revealed that there were very few cells experiencing detectable arrests and nuclear missegregations in the populations with the normal amount of telomerase, with 97.1% of mitoses classified as Normal Divisions (Figure 2B and S1B). In contrast, *est2p-17* and *est2p-30* mutants showed a great reduction in the frequency of normal divisions (28.6% and 54.2% Normal Divisions, respectively), suggesting that most mitoses in the cells with TI involved G2/M arrests prior a cell division. Although a larger fraction of those resulted in symmetrical nuclear divisions (Class 1), a striking increase in aberrant mitoses was also observed (Class 3). The majority of these mitoses belonged to the Class 3.2 events where nuclear divisions were not observed (Figure S1B). While observed frequencies of aberrant mitoses were at 11.5% and 8.8% of all the scored mitotic events in *est2p-17* and *est2p-30* populations respectively, the estimated probabilities of Class 3 events following a G2/M arrest were significantly higher. One can project that the Class 2 cells would eventually become either Class 1 or Class 3 mitotic events, proportionally to the frequencies already observed in a given experiment. Therefore, in the *est2p-30* mutants the probability of a cell to undergo an aberrant mitosis after the arrest calculated as a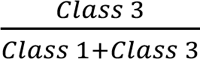 ratio was 0.32±0.06 or ∼32% (Figure S1C). Because *est2p-30* had an easily detectable aberrant mitosis phenotype while showing a milder growth defect in comparison with *est2p-17*, we used *est2p-30* for further experiments.

Following Class 3.1 and 3.2 aberrant mitoses, cells often died, most likely as a result of gross missegregation of nucleolus from the rest of the nucleus. We found that, indeed, the thread-shaped Htb2-GFP signal often remaining in the anuclear cell co-localised with the nucleolar protein Nop56 (Figure 2C-E). Despite inheriting most of the genome, the sibling cell eventually died (see Figure 2E-F for representative examples). We followed 654 *est2p-30* cells after either Class 3.1 or 3.2 mitosis. All of the nucleus-inherited cells budded and underwent another G2/M arrest in the following cell cycle (Figure 2G). Around 90% of them remained arrested by the end of imaging (Class 2). Of the remaining 75 cells, only 1 underwent a seemingly symmetrical nuclear division, 8 experienced another aberrant mitosis and 66 cells died. Therefore, telomerase insufficiency results in a vastly increased probability of aberrant mitoses and cell death.

### Aberrant mitoses result from DNA damage-induced G2/M arrests

Short telomeres are known to fuse through non-homologous end joining (NHEJ) which might then impair chromosome segregation (Mieczkowski et al. 2003). To test this hypothesis, we inactivated NHEJ by deleting *DNL4*, the gene coding for the DNA ligase responsible for telomere fusions. However, the loss of *DNL4* had no effect on chromosome segregation in cells with TI (Figure S2). Therefore, the increase in aberrant mitoses in response to TI was not due to telomere-telomere fusions.

We next asked if the residual telomerase in the *est2p-30* mutants played any role in the aberrant mitoses or this phenotype was due to their short telomeres. *EST2* was deleted in *HTB2-GFP* cells and the telomerase-deficient populations were assayed through the replicative senescence using time-lapse microscopy as described above. As the telomeres shortened (Figure S3A), the fraction of normal divisions decreased, reaching its lowest percentage at the point of the viability crisis (Figure S3B). At the same time, the fractions of the cells undergoing G2/M arrests increased and the aberrant mitoses paralleled this trajectory reaching the maximum frequency at the point of the cell viability crisis when the telomeres were the shortest. As the populations generated Type II survivors, the normal divisions became more frequent again and the aberrant mitoses decreased. Thus, the aberrant mitoses were not specific to the cells with TI, but correlated with the presence of short telomeres.

Telomerase-deficient cells undergo longer cell cycle arrests during the later stages of senescence (Xu et al. 2015). Therefore, the increase in aberrant mitoses during senescence observed in Figure S3 might be due to the increase in the duration of G2/M arrests. To test this hypothesis, we re-analysed the microscopy data on *est2p-30* mutants (Figure 2B) by binning the mitotic events according to the duration of the cell cycle. It was calculated as the time either between nuclear segregations in consecutive cell cycles in Normal Divisions and Class 1 events or between the preceding nuclear segregation and cutting off the thread-shaped nucleolar Htb2-GFP signal in Class 3 events. (Figure 3A-B). Indeed, the aberrant mitoses correlated with the longer cell cycles pointing toward the duration of the G2/M arrests being an important factor in determining the outcome of cell divisions.

**Figure 3.**
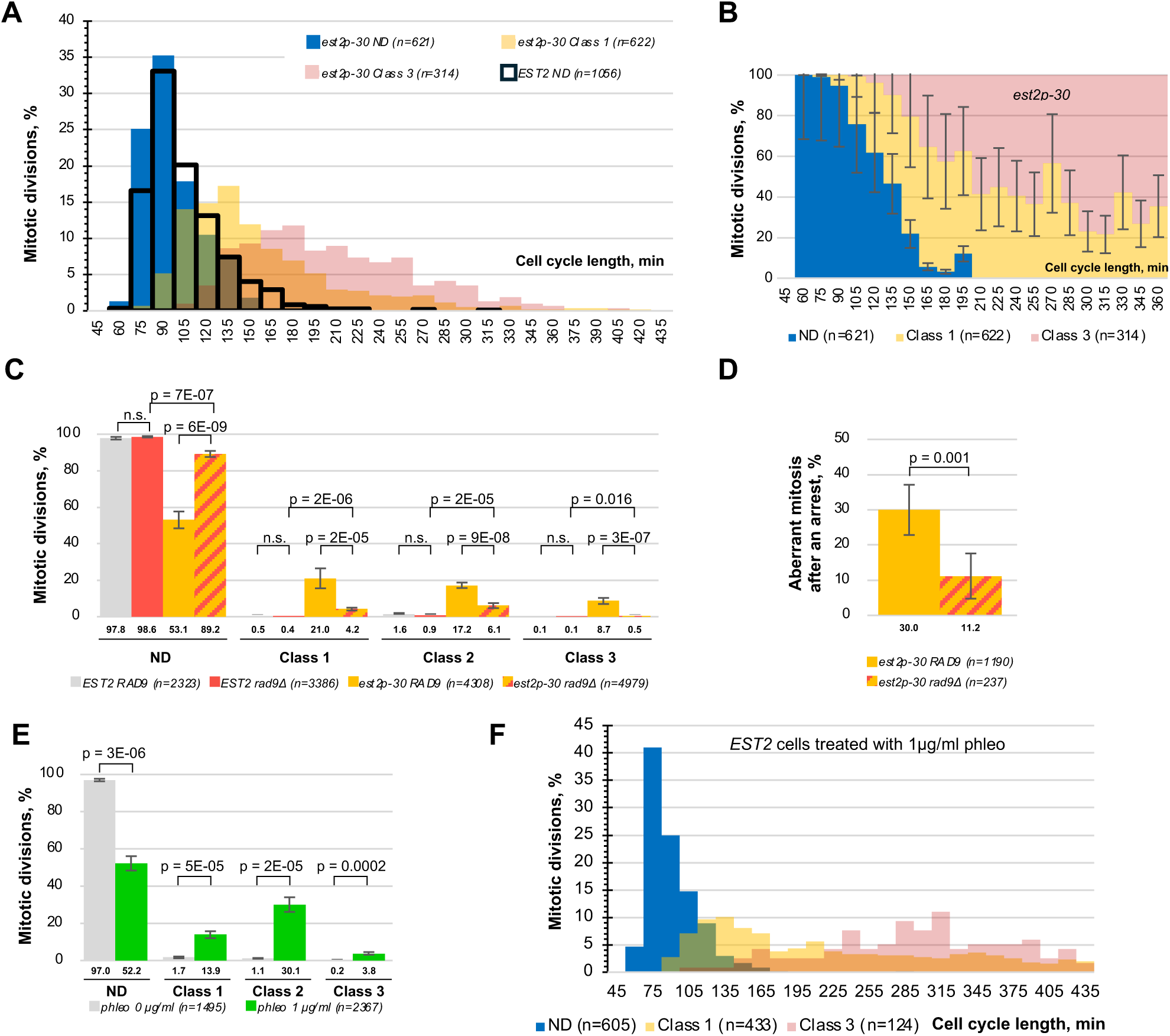
Aberrant mitoses result from DNA damage-induced G2/M arrests (A) Distribution of completed mitoses in *est2p-30* cells according to the cell cycle duration. (B) The probability of aberrant mitoses in *est2p-30* cells increases with the cell cycle length. (C) *RAD9* deletion suppresses high frequency of aberrant mitoses in response to TI. (D) Probability of aberrant mitoses in *est2p-30* cells undergoing G2/M arrest depends on *RAD9*. (E) Addition of phleomycin to the growth media results in an increase in aberrant mitoses. (F) Distribution of completed mitoses in wild-type cells grown in the presence of phleomycin according to the cell cycle duration, see panel E. (C-E) Averages (shown below each bar) of at least 3 biological repeats are plotted for each genotype. The error bars represent st.dev. A total number of mitotic events (n) scored for each genotype in all the repeats is stated in the legend.

Lack of the key component of the DNA damage checkpoint Rad9 in telomerase-negative cells leads to a dramatic reduction in the fraction of large G2 arrested cells (Sandell and Zakian 1993; Enomoto et al. 2002; AS and Greider 2003). The loss of *RAD9* in *est2p-30* cells resulted in a suppression of aberrant mitoses (Figure 3C). This suppression could not be attributed solely to the decrease of the fraction of the cells undergoing G2/M arrests, because when the frequency of aberrant mitoses was calculated only among the cells undergoing G2/M arrests (longer than 10 min G2), a significant decrease in aberrant mitoses remained (Figure 3D). Therefore, the DNA damage checkpoint activation increases the probability of an aberrant mitosis.

If the DNA damage-induced G2/M arrests contribute to aberrant mitoses, then a similar phenotype might be observed in response not only to short telomeres, but also to the DNA damaging drugs triggering the DNA damage signalling pathway. *HTB2-GFP EST2* cells were grown in the presence of phleomycin, a drug causing DNA breaks, and their mitotic behaviour was analysed using time-lapse microscopy (Figure 3E). In the presence of phleomycin, more cells underwent G2 arrests (lower NDs). More importantly, the fraction of aberrant mitoses also increased in response to phleomycin (Figure 3E, Class 3) and the aberrant mitoses correlated with the longer cell cycles (Figure 3F). Therefore, not only short telomeres, but also DNA damage in non-telomeric regions of the genome can cause genome missegregations. Taken together, all the experiments presented in Figures 3 and S3 suggest that prolonged G2/M arrests in response to DNA damage signalling triggered by either short telomeres or DSBs result in increased frequencies of aberrant mitoses.

### Accumulation of Pds1 during prolonged G2/M arrests leads to aberrant mitoses

What is the molecular mechanism linking the length of the G2/M arrest with the frequency of missegregation events? Strikingly, most of the aberrant mitoses are represented by the majority of the chromosomes being pulled all together in the same cell (Class 3.2), rather than random genome missegregation. This suggests the possibility that the sister chromatids remain connected. Pds1 (securin) is one of the downstream targets of the DNA damage signalling. Pds1 plays a key role in mitosis because it inhibits the sequence-specific protease Esp1 (separase). Separase cleaves cohesin at the onset of the metaphase-to-anaphase transition, allowing sister chromatin segregation. Pds1 is phosphorylated by the DNA damage checkpoint kinase Chk1 to stabilise Pds1 and this phosphorylation is required for prolonged G2/M arrests in response to DNA damage (Wang et al. 2001). We hypothesised that the stabilisation of Pds1 might lead to its accumulation with increasing lengths of the G2/M arrest and interfere with the activation of separase. First, to test if Pds1 phosphorylation played a role in the increase in aberrant mitoses in cells with TI, a previously characterised phosphorylation-deficient allele *pds1-m9* (nine serine/threonines are replaced with alanines (Wang et al. 2001)) was combined with *est2p-30* and the double mutants along with the appropriate isogenic control strains were analysed by time-lapse microscopy, as described above. The *pds1-m9* mutation did not affect the probability of a G2/M arrest in response to TI (Normal Divisions: 55.3% in *PDS1* vs. 60.1% in *pds1-m9*), but it drastically reduced the probability of an aberrant mitosis, from 6.2% in *PDS1* control to 0.1% in *pds1-m9* (Figure 4A-B). The reduction in the Class 3 events was accompanied by an increase in the Class 1 events, i.e. more post-arrest cells segregated their genomes with symmetrical nuclear divisions instead of going through aberrant mitoses. Similar results were obtained using phleomycin as a DNA damage signalling trigger instead of TI (Figure 4C). Therefore, the phosphorylation of Pds1 in response to DNA damage plays an important role in aberrant mitoses in cells with G2/M arrests.

**Figure 4.**
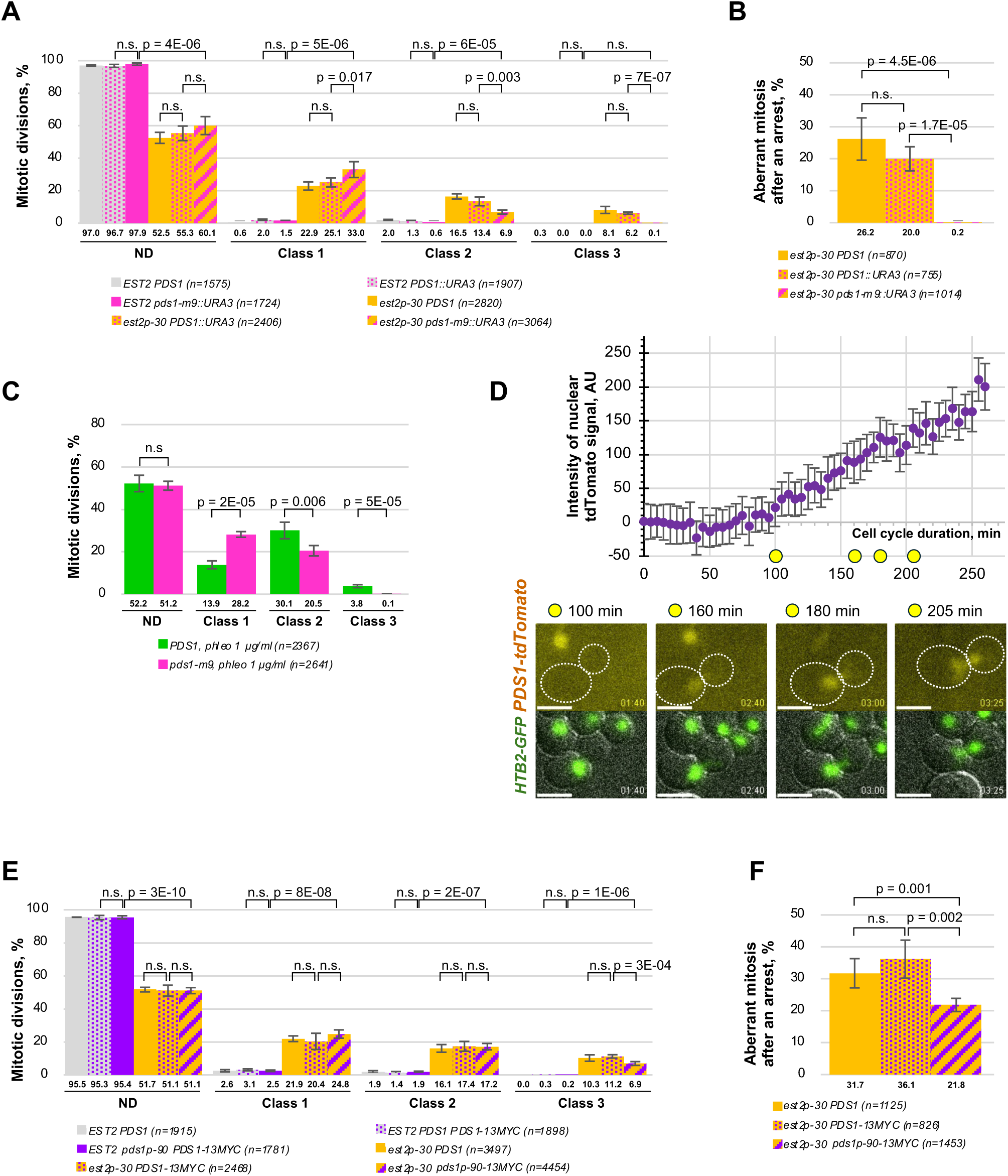
Accumulation of Pds1 during prolonged G2/M arrests leads to aberrant mitoses. (A) *pds1-m9* suppresses aberrant mitoses in *est2p-30* cells. (B) The probability of aberrant mitoses in *est2p-30* cells after undergoing G2/M arrest depends on *PDS1* phosphorylation. (C) *pds1-m9* suppresses aberrant mitoses caused by cell growth in the presence of phleomycin. (D) Analysis of Pds1-tdTomato accumulation during a G2/M arrest at the single-cell level. The intensity of nuclear tdTomato signal over background during the observed cell cycle is plotted. Cell images for 4 time points indicated by yellow circles on the X-axis are shown below the plot. 15 Class 2 events in *est2p-30* cells were analysed and all showed a similar Pds1-tdTomato dynamics to the presented example. Error bars show the standard deviation of pixel intensity. (E) *pds1p-90* partially suppresses aberrant mitoses in *est2p-30* cells. (F) Probability of aberrant mitoses in *est2p-30* cells undergoing G2/M arrest depends on *PDS1* expression levels. In panels A-C and E-F, averages of at least 3 biological repeats are plotted. The error bars represent st.dev. A total number of mitotic events (n) scored for each genotype in all the repeats is stated in the legend.

Next, we followed the levels of Pds1-tdTomato at the single-cell level in *est2p-30* mutants with prolonged G2/M arrests (Figure 4D). The Pds1-tdTomato signal was below the detectable levels at the earlier stages of the arrests but the protein gradually accumulated and became easily-detectable. Therefore, Pds1 progressively accumulates over the duration of G2/M arrests.

Finally, our hypothesis proposes that excessive accumulation of Pds1 during prolonged arrests might be responsible for aberrant mitoses by interfering with separase activation. Consequently, we reasoned that lowering the excessive Pds1 levels might suppress aberrant mitoses caused by TI. In order to decrease the *PDS1* expression in general, we used the same strategy as the one described above for *EST2* (Figure 1B-C). *kanMX6* cassette coding for a Kanamycin resistance gene was inserted at different positions in the *PDS1* promoter, in a strain with a *PDS1-13MYC* allele. The levels of Pds1-13MYC in the constructed mutants were analysed using anti-MYC antibodies (Figure S4A-B). To select the mutation that affected the accumulation of Pds1 upon activation, we grew the cells carrying a *kanMX6* insertion in different positions in the presence of phleomycin, to induce G2/M arrests. *pds1p-90* and *pds1p-50* mutants showed noticeable reductions in Pds1 levels. *pds1p-90-13MYC* was combined with *est2p-30* and the mitotic behaviour of the double mutant strains, as well as of the isogenic control yeast, was analysed using time-lapse microscopy (Figure 4E-F). Reducing the levels of *PDS1* expression did not decrease the detectable fraction of the cells experiencing G2/M arrests, but it led to a partial suppression of aberrant mitoses. Thus, excessive Pds1 in the cells with DNA damage induced G2/M arrests is responsible for more frequent aberrant mitoses.

### Aberrant mitoses stem from a loss of coordination between nuclear divisions and cytokinesis

In order to gain mechanistic insights into the aberrant mitoses, we co-visualised Htb2-GFP with one of the components of mitotic machinery, such as spindle microtubules (mRuby2-Tub1), spindle pole bodies (SPB, Spc42-tdTomato) and the actomyosin ring (Myo1-mCherry) involved in cytokinesis, and compared the dynamics of cellular components in *estp-30* cells during Class 1 and Class 3 mitoses (Figure 5). In both cases, the cells were dividing after a G2/M arrest, but underwent either seemingly normal or aberrant mitoses. In Class 1 cells, shorter spindles extended into long spindles characteristic of normal mitoses (Figure 5A) and the actomyosin ring contracted following chromosome segregation into mother and daughter cells (Figure 5B). In contrast, during Class 3.1 mitoses short spindles collapsed first and then re-established and elongated inside the buds/daughter cells during attempted nuclear divisions, which occurred within a single cell (Figure 5C). These nuclear divisions either reversed before the completion and resulted in a single undivided nucleus or went all the way to the end yielding binuclear cells.

**Figure 5.**
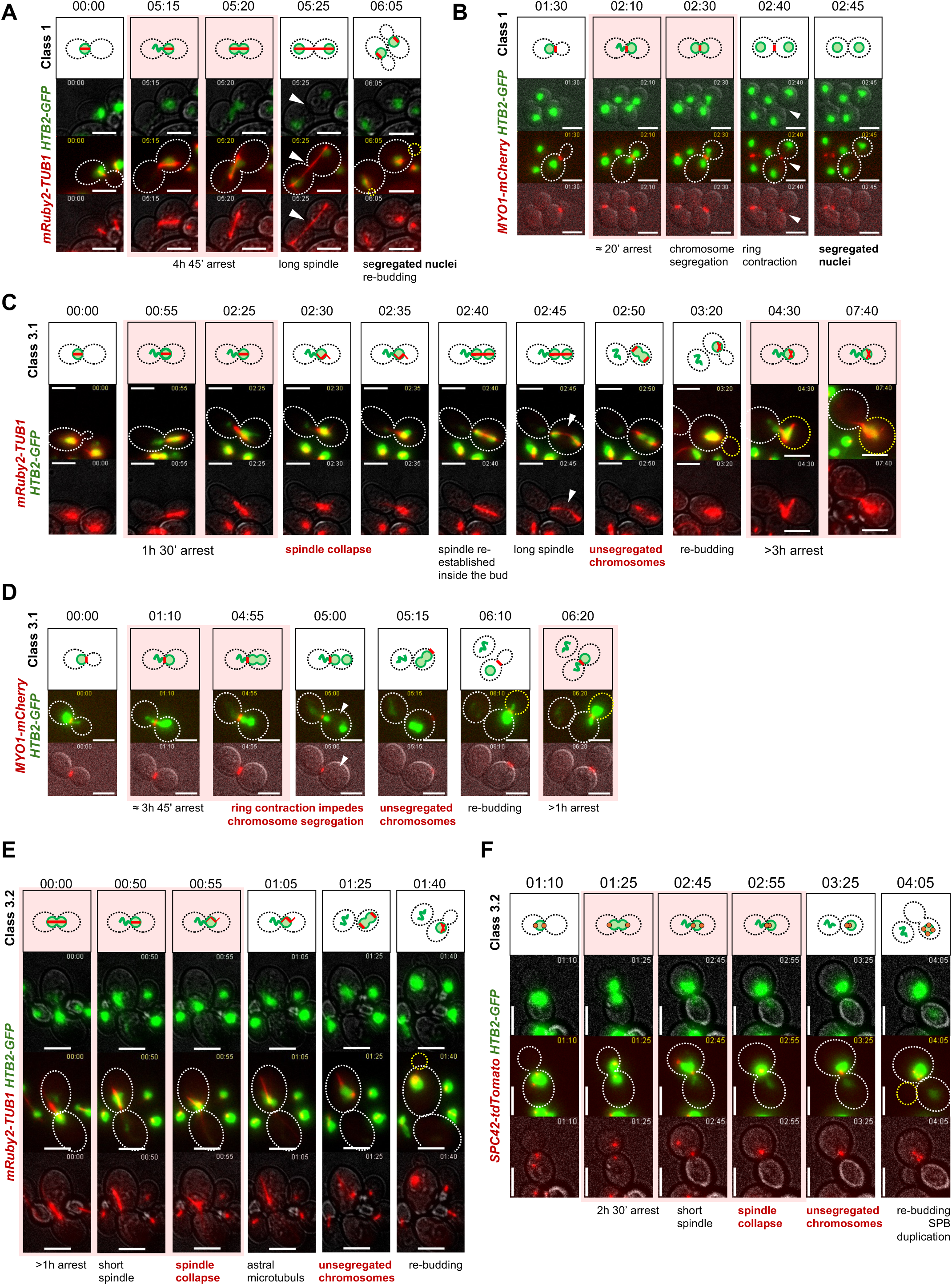
TI leads to a loss of coordination between nuclear division and cytokinesis. (A-F) Representative examples of mitotic events in *est2p-30* observed using time-lapse microscopy. Consecutive images of the same field are shown in each set, as explained in the legend to Figure 2. White arrows are pointing at nuclear divisions. White cell outlines (dotted lines) are used for original cells selected for analyses and yellow outlines show their buds and the progeny after re-budding. (A) Co-visualisation of mitotic spindle (mRuby2-Tub1) and chromatin (Htb2-GFP) during a Class 1 mitosis. (B) Co-visualisation of actomyosin ring (Myo1-mCherry) and chromatin (Htb2-GFP) during a Class 1 mitosis. (C) Co-visualisation of mitotic spindle (mRuby2-Tub1) and chromatin (Htb2-GFP) during a Class 3.1 aberrant mitosis. (D) Co-visualisation of actomyosin ring (Myo1-mCherry) and chromatin (Htb2-GFP) during a Class 3.1 aberrant mitosis. (E) Co-visualisation of mitotic spindle (mRuby2-Tub1) and chromatin (Htb2-GFP) during a Class 3.2 aberrant mitosis. (F) Co-visualisation of spindle pole bodies (Spc42-tdTomato) and chromatin (Htb2-GFP) during a Class 3.2 aberrant mitosis.

Why did the nuclear divisions happen inside the bud? We hypothesised that premature cytokinesis which occurs in yeast via actomyosin ring contraction at the bud neck might have created a physical barrier by separating the mother and daughter cells before the nuclear divisions took place. Indeed, in Class 3.1 divisions, the ring contraction began before the genetic material segregated into daughter cells (Figure 5D) and could be seen as an obstacle for the genome segregation.

During Class 3.2 mitotic events, short spindles collapsed, as evident by a) mRuby2-Tub1 signal converting into a single spot with radiating astral microtubules (Figure 5E) and b) merging SPBs (Figure 5F). As the nuclear division did not take place, the SPB re-duplication resulted in 4 bright Spc42-tdTomato spots within a single cell (Figure 5F). Taken together, these observations suggest that the aberrant mitoses might come from either delayed or completely missed nuclear divisions, where cytokinesis might happen before the nucleus could divide.

Based on the findings presented above, we propose a model where excessive accumulation of Pds1 during extended G2/M arrests results in aberrant cell divisions (Figure 6A). Upon metaphase-to-anaphase transition, Pds1 degradation is induced, but because of Pds1 accumulation during the arrests, it takes much longer than usual to reduce Pds1 to the levels low enough to activate the separase Esp1 and allow sister chromatids to separate. Meanwhile, the mitotic entry also leads to activation of the MEN pathway, which governs cytokinesis. Due to the delay in Pds1 removal, the cohesin cleavage and cytokinesis become mis-coordinated.

**Figure 6.**
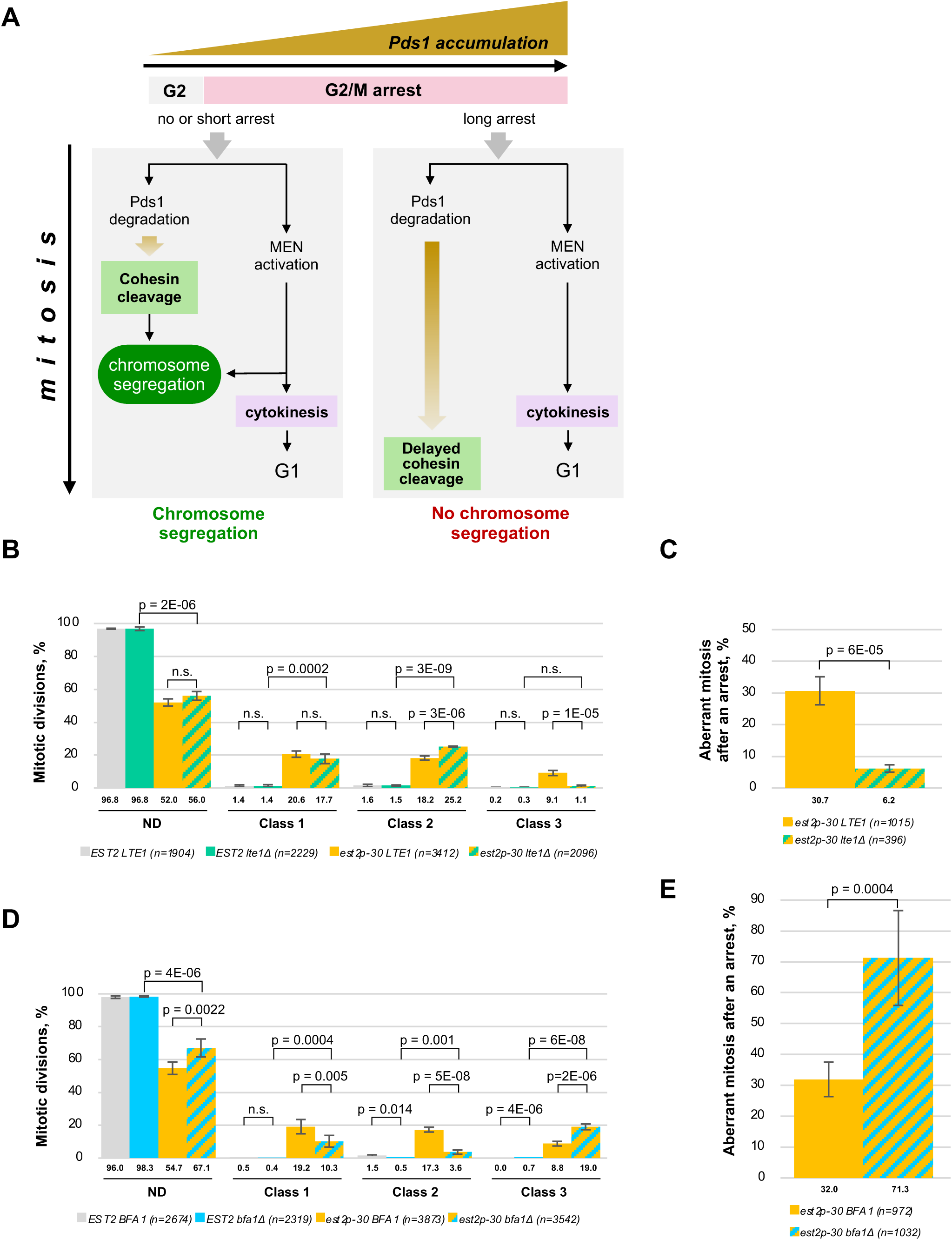
Activation of MEN affects aberrant mitoses. (A) Model for aberrant mitoses: mis-coordinated cohesin cleavage and MEN activation lead to skipped chromosome segregation and an aberrant mitosis (on the right). This miscoordination stems from accumulation of excessive Pds1 during extended G2/M arrests (top) which leads to a delay in completing the degradation of Pds1 and the inability of the cells to segregate chromosomes before cytokinesis takes place. (B) *LTE1* deletion suppresses high frequency of aberrant mitoses in response to TI. (C) Probability of aberrant mitoses in *est2p-30* cells undergoing G2/M arrest depends on *LTE1*. (D) *BFA1* deletion exacerbates high frequency of aberrant mitoses in response to TI. (E) Probability of aberrant mitoses in *est2p-30* cells undergoing G2/M arrest depends on *BFA1*. In panels B-E, averages of at least 3 biological repeats are plotted. The error bars represent st.dev. A total number of mitotic events (n) scored for each genotype in all the repeats is stated in the legend.

If this hypothesis is correct then manipulating the activation or progress of MEN may affect the probability of aberrant mitoses. In order to slow down MEN, we constructed TI strains lacking Lte1, an important, but not essential activator of the MEN cascade(Falk et al. 2016). Deletion of *LTE1* in the wild-type cells had no significant effect on the fractions of different mitotic events (Figure 6B), but the *est2p-30 lte1Δ* double mutants experienced significantly fewer aberrant mitoses than the single *est2p-30* mutants (Figure 6B-C). Also, the vast majority of these Class 3 events belonged to the Class 3.1 (0.8% from the total of 1.1% in class 3), i.e. a nuclear division was detected. This could be explained by shorter delays of the cohesin cleavage relative to cytokinesis in the *est2p-30 lte1Δ* mutants. These results are consistent with the loss of Lte1 delaying mitotic exit and therefore cytokinesis, thereby allowing more time for the degradation of excessive Pds1 and suppressing aberrant mitoses in cells with TI.

It has been reported that the inhibition of MEN through Bfa1 is essential to maintain the G2/M arrest in response to DNA damage signalling due to compromised telomeres (Valerio-Santiago et al. 2013). Consistent with the previous observations on premature MEN activation in cells with telomere damage, the *est2p-30 bfa1Δ* yeast were not able to maintain prolonged arrests (Figure S5A-C). More importantly, the frequency of aberrant mitoses in these populations was dramatically increased (Figure 6D-E), consistent with premature MEN activation promoting aberrant mitoses. Therefore, the cohesin cleavage and MEN activation are highly coordinated, but this coordination is at least partially lost during prolonged G2/M arrests due to excessive accumulation of securin.

### Reducing the levels of securin leads to an increase in aneuploidy

The reduction in Pds1 levels in *pds1p-90* mutants did not cause a detectable increase in Class 3 events in cells with wild-type telomerase (Figure 4E). Why haven’t yeast cells evolved a lower level of *PDS1* expression to avoid aberrant mitoses after prolonged G2/M arrests? One explanation is that it might have come at a higher cost of other types of genomic instability under normal growth conditions, for example, missegregation of single chromosomes due to a less tight control of cohesin cleavage. In order to address this hypothesis, we compared the rates of single chromosome mis-segregation in homozygous wild-type (*PDS1/PDS1*) and isogenic heterozygous (*PDS1/pds1Δ*) diploids. These experiments were performed in either *MAT a/α* or *MATΔ/α* background. The former is typical of normal diploid yeast, while the latter mimics haploids, where the homologous recombination pathway is significantly less active (Valencia et al. 2001). We constructed a genetic system, where the loss of one or the other copy of the chromosome V (*CHR V*) could be quantitatively assayed by first selecting for a marker loss using either 5FOA (loss of *URA3*) or canavanine (loss of *CAN1*), and then screening the post-selection colonies for the loss of another marker on the other arm of the same chromosome (Figure 7A-B). Lowering the Pds1 levels through heterozygosity resulted in an increase in the *CHR V* aneuploidy (Figure 7C). Analysis of the karyotypes of the identified aneuploids by PFGE confirmed the loss of *CHR V* in most analysed clones. The remaining clones might have re-gained a second copy of the remaining *CHR V* in the process of the experiment as returning to euploidy conferred a strong growth advantage. Furthermore, the loss of *CHR V* was often accompanied by either gains or losses of other chromosomes in many of these clones (Figure S6). Therefore, the Pds1 levels might have evolved to be optimal for providing accurate chromosome segregation under the normal growth conditions. In this case, the protein accumulation during the G2/M arrest allows the majority of cell divisions to occur normally. It remains to be addressed whether the observed death of the long-arrested cells through aberrant mitoses is collateral damage or an evolutionary product designed to eliminate the cells struggling to repair DNA damage. These cells are the most likely candidates for undergoing genomic instability should they manage to survive DNA damage stress and eliminating them through aberrant mitoses might be a quality control mechanism preserving genomic stability in microbial populations.

**Figure 7.**
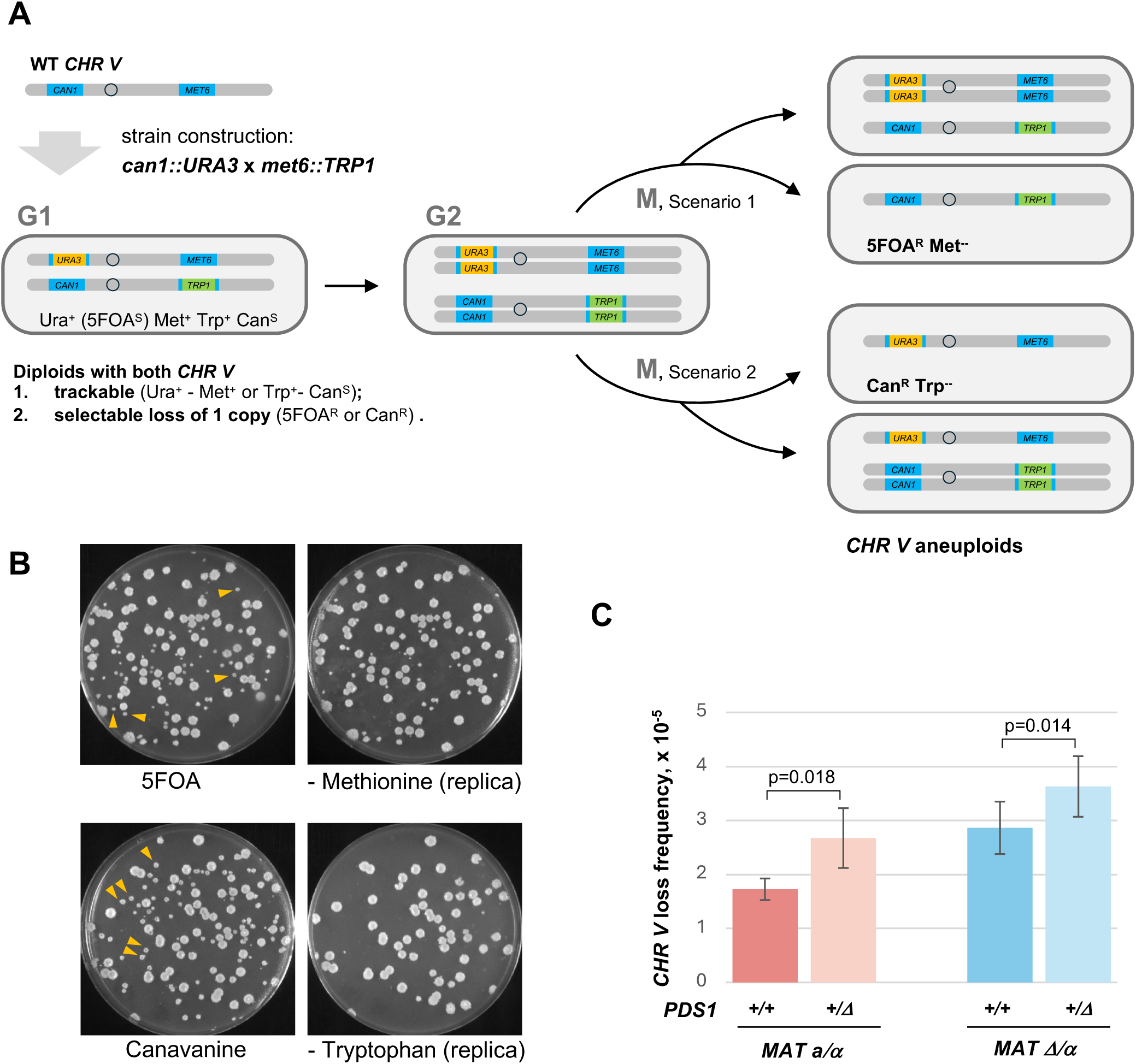
*PDS1* heterozygosity results in an increase in aneuploidy (A) Schematic of the experimental setup for quantitative analysis of single chromosome loss in diploid cells. Both copies of *CHR V* were marked with two different genetic markers, one on each arm, in haploid cells of opposite mating type. In post-mating diploids, one copy of *CHR V* contained *URA3* and *MET6,* while the other had *CAN1* and *TRP1* at the exact same locations as *URA3* and *MET6* respectively. A loss of *CHR V* with *URA3-MET6* due to mis-segregation (Scenario 1 on the right) results in cells which can be selected on 5FOA and screened for Met^-^ on plates without methionine. Similarly, a loss of the *CAN1-TRP1* copy of *CHR V* (Scenario 2) leads to Canavanine resistance and a Trp^-^ phenotype. (B) Representative examples of *CHR V* loss analysis using selection for either 5FOA-resistant or Canavanine-resistant colonies. The former were screened by replica-plating for Met^-^ (top pair of plates) while the latter for Trp^-^ (bottom pair of plates). Notice that *CHR V* aneuploids have a growth defect and appear as small colonies, some of which are indicated with yellow arrowheads. (C) *PDS1* heterozygocity results in an increase in *CHR V* loss. Averages of at least 3 biological repeats are plotted for each genotype. The error bars represent st.dev.

## DISCUSSION

In this study, we described at the single cell level how activation of DNA damage signalling in response to DSBs and critically short telomeres affects the cell cycle progression, in particular genome segregation in mitosis. In order to investigate cells with critically short telomeres, we constructed a yeast model for TI. In many aspects, this model mimics the developmentally programmed TI in humans and other large mammals where the gene for telomerase catalytic subunit TERT is downregulated during development (Wright et al. 1996; Gomes et al. 2011). In contrast, yeast express telomerase constitutively, but TI in *S. cerevisiae* can be triggered naturally by higher growth temperature, which leads to lower Est2 protein levels (Millet et al. 2015). Here, we decreased Est2 by interfering with the *EST2* gene promoter and generated cell populations with TI under the growth conditions normally used in laboratory settings.

Cells with TI are different from senescing telomerase-deficient mutants: telomerase activity is still present and given enough time, all the critically short telomeres can be extended to allow inactivation of the DNA damage signalling and cell cycle progression. Telomerase-deficient cells have no means to repair their problem telomeres other than through recombination, which can only be used on Y’-telomeres due to their shared homology in the sub-telomeric regions. The time window for telomerase to operate in cells with TI can be extended by a G2/M arrest triggered by critically short telomeres. DNA damage signalling also stimulates homologous recombination (Bashkirov et al. 2000; Barlow and Rothstein 2009; Flott et al. 2011; Ullal et al. 2011; Dion et al. 2012). Recombination might be either regularly or occasionally operating on Y’-telomeres in cells with TI, as evident by the amplification of sub-telomeric Y’ repeats (Figure 1D) characteristic of Type I telomerase-deficient survivors (Lundblad and Blackburn 1993). Because all Y’-telomeres share extended homology, Rad51-dependent recombination can be used as a mean to repair one Y’-telomere by using another one as a DNA template donor (Louis and Haber 1990; Lundblad and Blackburn 1993; Le et al. 1999; Chen et al. 2001). In Type I survivors, X-telomeres also acquire Y’-repeats to join the pool of the telomeres maintained via recombination. In contrast, recombination in cells with TI might be limited to the Y’-telomeres because the X-telomeres maintain the original structure without Y’ repeats (Figure 1D). However, the involvement of Rad51-dependent recombination in the Y’-telomere maintenance might ease the load for the low-expressed telomerase and improve its availability for X-telomeres in the cells with TI. Therefore, the cells with TI might be using both Rad51-dependent recombination and telomerase for telomere maintenance.

A single unrepairable DSB leads to a G2 arrest which extends the cell cycle duration approximately 6 times (Dotiwala et al. 2007). Following this long arrest, cells undergo Cdc5-dependent adaptation by dividing in spite of the unrepaired damage (Sandell and Zakian 1993; Toczyski et al. 1997; Dotiwala et al. 2007). Telomerase-deficient cells also involve adaptation at the later stages of senescence. The post-adaptation mitoses in these cells are often non-terminal, i.e. followed by further cell proliferation (Coutelier et al. 2018). In the cells with TI, we very rarely saw cell cycle duration exceeding 360 minutes, i.e. only 4 normal cell cycles of 90 minutes each (Figure 3A). The distribution of aberrant mitoses peaked at around 180 minutes (Figure 3A), i.e. the duration of only 2 cell cycles, which might be too early to activate adaptation. Therefore, the aberrant mitoses described here might be different from the adaptation-driven divisions in telomerase-deficient cells (Coutelier et al. 2018). Instead of involving adaptation mechanisms, the cells with TI might rely on shutting down DNA damage signalling through successfully extending their critically short telomeres. However, the longer it takes to finish the repair the higher the probability of an aberrant mitosis becomes, as Pds1 gradually accumulates over the time of a G2/M arrest (Figure 4D). The terminal nature of aberrant mitoses might be responsible for cell death during senescence of telomerase-negative cells, when there is no telomerase and cells accumulate more and more uncapped telomeres as they senesce. Consistent with this idea, aberrant mitoses progressively increase with the bulk telomere shortening, peaking exactly at the cell viability crisis (Figure S3), but it remained unclear if adaptation plays a role in this process.

Mitosis is one of the most sophisticated stages of the cell cycle. It is orchestrated by multiple kinases and phosphatases, as well as the anaphase-promoting complex (APC) targeting proteins for degradation. The regulation of Pds1 is known to involve Cdk1 and Chk1 kinases, PP2A^Cdc55^ phosphatase and APC-directed proteasomal degradation upon cell commitment to mitosis (Wang et al. 2001; Agarwal et al. 2003; Khondker et al. 2020). Pds1 degradation triggers nuclear division, which is coordinated with cytokinesis controlled by MEN. The coordination is carried out by Cdc14 phosphatase and a polo-like kinase Cdc5 (Cohen-Fix et al. 1996; Shou et al. 2002; Stegmeier et al. 2002; Manzano-Lopez and Monje-Casas 2020). Activation of the DNA damage signalling, which arrests the G2/M progression, adds another level of regulation to this already complicated network. Chk1 phosphorylates Pds1 to stabilise it and further reinforce the block on chromosome segregation (Cohen-Fix and Koshland 1997; Sanchez et al. 1999; Wang et al. 2001). The mitotic progression after longer G2/M arrests does not always adequately adjust to the excess of accumulated Pds1 (Figure 5). Shorter delays in Pds1 removal may result in the Class 3.1 mitoses, where attempted nuclear divisions can be observed, even though they occur when the actomyosin ring is already closing or just closed. As a result, the nuclear division takes place inside one of the daughter cells due to the physical barrier at the bud neck. The cells with the Class 3.2 aberrant mitoses do not attempt nuclear divisions, possibly because by the time Pds1 is removed and the cohesin cleavage is complete, the cells are in the following G1 and have an inappropriate cyclin environment for the anaphase to take place. Class 3.3 mitoses, characterised by multipolar spindles, might stem from Class 3.1 or 3.2 divisions in the previous cell cycle, when SPBs do not segregate correctly (Figure 5F). Consistent with this hypothesis, Class 3.3 nuclear divisions were observed in the cells, which underwent aberrant mitoses but escaped cell death in the next cell cycle (Figure 2G). The hypothesis of miscoordination between nuclear division and cytokinesis being the primary reason for these aberrant mitoses (Figure 6A) is supported by our genetic manipulations of MEN progression. Delaying mitotic exit, by using *LTE1* deletion, partially suppresses aberrant mitoses, while its premature activation in *bfa1Δ* further exaggerates the phenotype (Figure 6B-E).

The aberrant mitoses in cells with TI and in response to phleomycin treatment resemble the terminal genome-level missegregation (GLM) events observed in ageing mother cells in *S. cerevisiae* (Crane et al. 2019). Like Class 3 events here, they are Rad9-dependent and characterised by the rDNA region on yeast chromosome XII remaining in a mother cell while the rest of the nucleus is moved to the bud. Yeast cells with terminal GLMs lack Cdc14 release from the nucleolus (Crane et al. 2019). During mitosis, Cdc14 is released in two stages. Although the early Esp1-induced partial release of Cdc14 can be dispensable for mitotic exit (Lu and Cross 2009), it is required for nuclear positioning, rDNA segregation, spindle stability and action prior to mitotic exit (D’Amours et al. 2004; Ross and Cohen-Fix 2004; Sullivan et al. 2004; Higuchi and Uhlmann 2005). Accumulation of Pds1 during prolonged G2/M arrests in cells with TI, DSBs and possibly in ageing mother cells would prolong Esp1 inhibition and delay or even abolish the Esp1-dependent release of Cdc14 from the nucleolus and might result in a mitotic exit accompanied by a genome missegregation. Therefore, the mis-regulation of the cohesin-independent function of Esp1 might contribute to the cause of aberrant mitoses.

Class 3 mitotic events lead to cell death. This cell death might result from losing rDNA and/or some essential genes in the close proximity to the rDNA array on chromosome XII in yeast. The aberrant mitoses seem to stem from a contracting myosin ring cutting off the nucleolus from the rest of the nucleus. While a 10-fold reduction in the number of the rDNA repeats, from ∼150 copies normally present in yeast down to 15 is compatible with cell survival (Kobayashi 2006; Iida and Kobayashi 2019), loss of the whole rDNA array would be lethal. The residual Nop56-mCherry signal is visible in some nuclei after aberrant mitoses (Figure 2E), but not in others (Figure 2D). However, the inheritance of residual rDNA array does not seem to correlate with the terminal fate of the cells, which might be also be missing other essential genetic material or dealing with unrepairable DNA damage. The disappearance of Htb2-GFP signal prior to cell death (Figure 2E-F) can be explained by degradation of histones previously observed in aged mother cells and in response to DNA damage (Hauer et al. 2017; Crane et al. 2019). Another explanation for the cell death might be activation of a pathway where Pds1 accumulation might be only one of the steps in a more complex mechanism, for example apoptosis. Interestingly, Pds1 and Esp1 have been linked to the regulation of H_2_O_2_-induced apoptosis in yeast (Yang et al. 2008), but it is unclear if this mechanism might be relevant to the cell death described in this study. More research would be needed to gain further molecular insights into how cells die following aberrant mitoses.

The DNA damage response functions to suppress genomic instability by allowing cells to pause the cell cycle until the repair is complete. This mechanism has an important additional level of regulation in multicellular eukaryotes, where the inability to complete repair or deal with other stresses in a timely manner can trigger one of the two developmental programs, either apoptosis or senescence, in order to eliminate cells with an increased potential for instability (reviewed in (Childs et al. 2014; Childs et al. 2017)). Our experiments suggest that unicellular organisms might use a similar strategy to preserve genome integrity in their populations. The probability of an aberrant mitosis becomes over 50% if a G2 arrest extends the cell cycle more than 200 min and further increases at even longer arrests (Figure 3B). Such a high probability of aberrant mitoses eventually leading to cell death would be an efficient enough mechanism to eliminate most cells struggling with DNA repair and therefore being at a higher risk of acquiring mutations. Even though these mutations could result in a growth defect in haploids, thereby limiting their spreading in the populations, mating to a healthy growing cell might allow the mutations to persist in a heterozygous state. Therefore, a mechanism involving Pds1 accumulation might have evolved to safeguard genome stability in the populations of unicellular eukaryotes.

Pds1 might operate as a stopwatch similar to the recently discovered p53-53BP1-USP28 mitotic stopwatch in human cells, which works via stabilisation of p53 (Meitinger et al. 2024). The p53-53BP1-USP28 complex “monitors” the duration of M-phase and triggers irreversible G1 arrests after prolonged mitoses, thereby protecting against defective mitotic events, which might generate potentially dangerous cells with unstable genomes. The Pds1-dependent pathway might be similar to the p53 mitotic stopwatch in how these two mechanisms are built – a sensor protein, Pds1 in yeast or p53 in mammals, is stabilised, accumulates over time and its levels then are used by the cell as a readout of time. The biological roles of these pathways are even more similar: they both eliminate potentially unstable cells to preserve genome integrity.

Lowering the *PDS1* gene expression levels partially suppresses the aberrant mitoses after prolonged G2/M arrests (Figure 4E-F), but at a cost of aneuploidy under the normal growth conditions (Figure 7C). These data argue that the evolution might have kept the Pds1 levels high enough to maximise the genome stability in unchallenged cells and that the Pds1 levels are finely tuned to minimise possible aneuploidy in the absence of stress, and then provide enough range during the G2/M arrests to differentiate between the cells with shorter and longer arrests.

Despite the very frequent cell death following the Class 3 events, there is a small chance for a cell to survive an aberrant mitosis and preserve its proliferation potential (Figure 2G, Class 1). Such cells would end up undergoing diploidisation, similar to tetraploidisation in higher eukaryotes following a mitotic slippage. Among other possible causes, compromised telomeres in p53-deficient human cells can lead to tetraploidisation in laboratory conditions (Davoli et al. 2010) and likely during carcinogenesis. Polyploidisation increases cells’ ability to adapt through aneuploidy by allowing them to reduce the number of some particular chromosomes to re-balance the genome to suit a particular adaptation. This is commonly found in cancers (Gordon et al. 2012) and has been also observed in yeast (Chen et al. 2012; Millet et al. 2015). Therefore, surviving mitotic slippage could be advantageous for adaptation of microbial populations to some stresses, and might play an important role in microbial pathogenesis where ploidy change and aneuploidy are very common (Todd et al. 2017; Gilchrist and Stelkens 2019). Understanding the genome dynamics in pathogenic fungi and protozoa in general and as a response to therapeutic treatments in medicine and crop protection measures in agriculture has a vast practical importance in improving human health and wellbeing. In achieving this, the single-cell approach allows one to account for the stochasticity factor in modelling the behaviour of cell populations without overlooking infrequent events, such as aberrant mitosis, as often they are the ones with the highest impact on how populations adapt to stress and evolve over time.

## MATERIALS AND METHODS

### Yeast strains, oligonucleotides and plasmids

Yeast strains, oligonucleotides and plasmids used in this study are described in Supplemental Tables S1, S2 and S3, respectively.

### Population doubling time measurement

Cultures in YPD medium were prepared by resuspending cells from freshly grown patches and grown overnight (at least 8 hours) to log-phase (OD_600_ < 0.6). The log-phase cultures were diluted using pre-warmed YPD broth to OD_600_ of 0.025. 200 µl of the resultant culture was transferred into a well of a sterile 96-well plate. Every analysed genotype had at least two biological and two technical repeats. The optical density was measured in a Tecan Infinite M200 plate reader. The absorbance curves were analysed using the Omniplate software (Montano-Gutierrez et al. 2022).

### Telomere analysis by Southern blotting

Telomere length and structure were analysed by telomere-specific Southern blotting. Purified yeast genomic DNA was digested with XhoI, resolved on 0.7% agarose gel in 1×TBE and transferred onto a nylon membrane. The membrane was hybridized to a ^32^P-radiolabeled telomeric oligo (CCCACA)_4_ to detect internal Y’ repeats and terminal restriction fragments.

### Western Blotting

Protein extracts for Est2-13MYC, Pds1-13MYC and actin western blotting were resolved in SDS polyacrylamide gel and transferred onto PVDF transfer membranes (Immobilon®-FL, 0.45 μm pores, Merck Millipore Ltd., IPFL00005, Darmstadt, Germany). Anti-c-Myc (9E10) mouse monoclonal (Thermo Fisher Scientific, Loughborough, UK, (13-2500), 1:1,000 dilution) was used to detect Est2-13Myc and Pds1-13Myc. Anti-β-actin mouse monoclonal antibody (Abcam ab8224, Cambridge, UK, 1:50,000 dilution) was used for Act1 western blotting. Goat anti-mouse IgG (H + L) cross-adsorbed secondary antibody Alexa Fluor 680 (Thermo Fisher Scientific, Loughborough, UK, A-21057, 1:12,500 dilution) was used with the corresponding primary antibodies. Western blotting membranes were scanned using Odyssey® CLx fluorescent scanner (LI-COR®, Cambridge, UK). The resulting images were analysed using Image Studio™ Lite software.

### Time-lapse and fluorescence microscopy

Imaging of live cells was carried out using a NikonTi2 inverted microscope equipped with a 100X 1.49 NA CFI Plan Apochromat TIRF objective, Lumencor Spectra X light source (Lumencor, Beaverton, OR USA) and a Photometrics Prime 95B camera (Teledyne Photometrics, Birmingham, UK). Filter sets from Semrock (Semrock, Rochester, NewYork, USA) were used to image GFP at excitation 474/26 nm, emission 525/40 nm, mRuby2 and tdTomato at excitation 543/23 nm, emission 595/31 nm and mCherry at excitation 578/21 nm, emission 641/75 nm.

In experiments to analyse the frequency of different mitotic events using chambers, yeast cultures were pre-grown in SC complete medium for at least 8 hours so that their OD_600_ did not exceed 0.6. Approximately 0.02 OD of the cell culture was added by pipetting into each well of the Ibidi chamber, pre-coated with concanavalin A. The images were taken every 5 min for at least 4 hours in the Brightfield and GFP channels with 5 Z-stacks of 0.6 μm. The GFP channel was set to 30 ms exposure time and 5% light source intensity.

We used agar pads to analyse cell area distribution. Agar pads were made by moulding the YPD liquid medium with 1% agarose in 65 µl Gene Frames. Yeast pre-grown in YPD were spotted on an agar pad and the sample was sealed with a coverslip. Multiple photos were taken in the Brightfield and GFP channels with 5 Z-stacks of 0.6 μm.

We also used agar pads to analyse the mitotic behaviour of cells with Myo1-mCherry, Spc42-tdTomato, Nop56-mCherry, mRuby2-Tub1, and Pds1-tdTomato. The images were taken every 5 min for at least 7 hours in the Brightfield, GFP and mCherry/tdTomato channels with 5 or 7 Z-stacks of 0.6 μm (Table S4).

Agar pads were also used to analyse the mitotic behaviour of cells treated with phleomycin. Cell cultures were grown in YPD at 30°C in a shaking incubator for at least for 8 hours until the OD_600_ reached 0.3. Phleomycin was added to the cultures at the desired concentrations and the cultures were left in the shaking incubator for another hour before being spotted on agar pads. The agar pads contained the same concentration of phleomycin as the treatment in the liquid culture. The images were taken every 5 min for at least 7 hours in the Brightfield and GFP channels with 5 Z-stacks of 0.6 μm. The GFP channel was set to 30 ms exposure time and 5% light source intensity.

### Image analysis and single-cell lineage tracking

The images were analysed using Fiji (ImageJ) software. Cell area was measured for all cells with clear borders in the focal plane using the polygon selection tool and excluding the bud cell compartment.

In time-lapse imaging, the mitotic behaviour of each cell in the focus plane was observed and categorised. A cell was considered undergoing an arrest if the majority of the DNA was transferred and localised in the bud for more than a single time frame (5 min). After that, the observed cell division with the arrest was assigned to Classes 1, 2 or 3, depending on the division outcome. The divisions completed after an arrest without any visible defect were defined as Class 1. Class 3 events were aberrant mitoses with an obvious defect in nuclear segregation. We observed splits of chromatin masses (transient or permanent) inside either a mother or a daughter cell compartment followed by re-budding of the cell with the majority of the DNA. These divisions were called Class 3.1 events. There were also cells generating an anucleate cell and a re-budding cell with the majority of the DNA without a prior nuclear division detected. These were called Class 3.2 events. Any division generating more than two separated chromatin masses was assigned to Class 3.3 events. Cells remaining arrested by the end of the recorded imaging were registered as Class 2 events.

Fiji (ImageJ) software was used for quantitative analysis of Pds1-tdTomato levels throughout the cell cycle. The mean fluorescence intensity and standard deviation of the tdTomato signal were measured within the nucleus, which was defined by Htb2-GFP signal. The background signal of the non-fluorescent area outside the cell was subtracted from the mean fluorescence intensity.

### Computational and statistical analyses

All statistical analyses in this study were performed using Student’s two-sample two-tailed t-test.

### Quantitative analysis of *CHR V* loss

Diploid strains with both armes of *CHRV* marked genetically *CAN1_met6::TRP1/can1::URA3_MET6 ura3-52/ura3-52 trp1-289/trp1-289* were streaked to single colonies on synthetic agar media without uracil and tryptophan. Single colonies grown in 48 h were patched on YPD agar and incubated for 24 h at 30°C. Patches were resuspended in water and appropriate serial dilutions were plated on YPD plates to determine the cell titre, on plates with 5FOA to select for colonies with lost *URA3*, and on plates with canavanine to select for a loss of *CAN1*. Colonies on YPD plates were scored in 2 days. In 3 days after the platings, colonies grown on the agar with the drugs were replica-plated from 5FOA onto plates without methionine and from canavanine onto plates without tryptophane. 5FOA^R^ Met^-^ and Can^R^ Trp^-^ colonies were scored as aneuploid. The frequency of aneuploidy was calculated as the ratio of aneuploids (5FOA^R^ Met^-^ and Can^R^ Trp^-^ combined) to the cell titre.

### Pulse-field gel electrophoresis of yeast chromosomes

The karyotypes of the cells selected for the loss of *CHR V* using the genetic markers were analysed by pulse-field gel electrophoresis as described previously (Hage and Houseley 2013). Colonies from the selective plates (5-FOA or CAN) were purified on YPD plates. A smaller colony characteristic of aneuploid phenotype was selected from each purification and patched on YPD overnight. A patch of a control WT haploid strain was included in each set of samples analysed on the same gel. Next day, the cells were washed with 1 ml of Wash Buffer WB (10mM Tris-HCL pH7.6, 50 mM EDTA) and re-suspended in 100 µl of WB with 17 U of Lyticase (Sigma L2524). The suspension was mixed with 100 µl of 1.6% agarose (Seakem® LE) pre-warmed to 50°C, and transferred into the plug mould. Agarose plugs were incubated with 500 µl of the Lyticase solution (170 U of Lyticase in 500 µl of WB) for 1 h at 37°C, and then with 500 µl of Proteinase K solution (100 mM EDTA, 0.2% SDS, 1% N-laurylsorcosine sodium, 1 mg/ml Proteinase K) overnight at 50°C. Next day, the plugs were washed with 1 ml of WB for 30 min at room temperature with gentle agitation, the wash was repeated three times. Half of each plug and the Yeast Chromosomes PFG marker (N0345S) were attached to a comb and placed into the gel casting tray. 1% agarose gel (Certified Megabase Agarose, Bio-Rad) was poured over the plugs and polymerised at 4°C, the comb was gently extracted. Chromosomes were resolved using CHEF-DR III electrophoresis system (Bio-Rad) in 0.5x TBE at 6 V/cm and 14°C with the following current switch settings: 120° every 60 sec for 16 h, followed by 120° every 90 sec for 11 h. After the run was finished, the gel was stained with ethidium bromide and imaged using Gel Doc XR+ imaging system (BioRad).

For the analyses of aneuploidy (only *CHR X, XI, V, VIII, IX, III, VI* and *I*) the signal in the band of each chromosome in the gel was normalised against the same chromosome in the control haploid strain and the resultant values for the eight analysed chromosomes were used to calculate an average for the normalised values for each strain. Then, the normalised value for each chromosome was divided by this average to calculate the relative increase/decrease in the EthBr staining for each chromosome. The values often deviated from a clear indication of a chromosome gain or loss, perhaps, because the populations of aneuploid cells are highly unstable Reverting back to euploidy often confers a growth advantage and therefore at least some analysed clones might be mixed populations of cells with different karyotypes.

## DECLARATION OF INTERESTS

The authors declare no competing interests.

## ACKNOWLEDGMENTS

We thank Sara Buonomo and Kevin Hardwick for critical reading of the manuscript, Sara Buonomo, Kevin Hardwick, Adele Marston, Meriem El Karoui, Peter Swain and the members of the Makovets lab for advice and helpful discussions, Peter Swain, Ivan Clark and Dave Kelly for help with setting up the imaging technique, Yu Huo for help with setting up the plate reader measurements, Eric Wooten and Stephen Elledge for the *pds1-m9* plasmid, John Hutchinson for constructing initial mutants with TI, and the Darwin Trust for a PhD Fellowship to A.D. This work was supported by funding for the Wellcome Discovery Research Platform for Hidden Cell Biology [226791] and we gratefully acknowledge support from the Light Microscopy core. The research was funded by MRC Senior Non-Clinical Fellowship to S. M., grant number MR/R02068X/1.

## AUTHOR CONTRIBUTIONS

A.D and S.M discussed the project and developed the concept. A.D. did the experiments in Figures 1-6 and S1-S5. M.J. and S.M. did the experiments in Figure 7 and O.K. performed the PFGE in Figure S6. A.D. and S.M. made all the figures and S.M. wrote the manuscript, with the inputs from all the authors.

